# β-helical protein PerA orchestrates antibiotic uptake across the *Caulobacter* outer membrane via converging stress signaling systems

**DOI:** 10.1101/2023.03.30.534887

**Authors:** Jordan Costafrolaz, Gaël Panis, Simon-Ulysse Vallet, Manuel Velasco Gomariz, Laurence Degeorges, Kathrin S. Fröhlich, Patrick H. Viollier

**Author notes:** corresponding author: Patrick H. Viollier.

## Abstract

While large membrane-impermeable antibiotics cannot traverse a lipid barrier, spacious importers such as TonB-dependent receptors (TBDRs) can mistakenly ferry antibiotics across the bacterial outer membrane (OM). We discovered that loss of PerA, an enigmatic β-helix protein in the OM of the oligotrophic α-proteobacterium *Caulobacter crescentus*, reprograms the OM TBDR proteome from ChvT that imports the glycopeptide antibiotic vancomycin to an uncharacterized TBDR (BugA) that confers sensitivity to the polypeptide antibiotic bacitracin. Both antibiotics are large zinc-binding molecules that target the peptidoglycan, echoing the zinc stress response induces destabilization of PerA. Inactivation of PerA launches two conserved and interwoven envelope stress programs that remodel the OM with TBDRs, a tripartite multidrug efflux pump and periplasmic proteases. Thus, unanticipated entry routes for antibiotics emerge in stressed diderm bacteria that may be treatable with membrane impermeable antimicrobials owing to an underlying transcriptional stress response pathways coordinated by a novel type of OM regulator.

## INTRODUCTION

The outer membrane (OM) protects diderm bacteria from toxic compounds by obstructing their entry into the periplasmic space where the essential cell wall (peptidodglycan layer), a major antibiotic target, is located (Sun et al., 2022). While the cytoplasmic membrane is a phospholipid bilayer, the OM features an inner leaflet of phospholipids and an outer leaflet with the charged glycolipid lipopolysaccharide (LPS). The OM barrier function also comes at a price, since essential nutrients can only traverse the OM through pores formed by β-barrel proteins. While small solutes pass through ungated pores, larger nutrients typically depend on active uptake mediated by gated channels such as TonB-dependent receptors (TBDR) that can open or close using the energy transmitted from the IM proton motive force via the ExbBD-TonB complex. TBDRs are β-barrel proteins known to import diverse substrates such as siderophores or metallophores (Moeck and Coulton, 1998; Pawelek et al., 2006), ionophores (Schauer et al., 2007), vitamins (Shultis et al., 2006), lignin derivatives (Fujita et al., 2019) and carbohydrates (Blanvillain et al., 2007; Neugebauer et al., 2005), but they can also serve as receptors for phages and antimicrobial peptides such as colicins (Braun et al., 1976; Salomon and Farias, 1993). Ungated TBDRs are wide-enough to allow the passage of large soluble antibiotics(Killmann et al., 1993; Krishnamoorthy et al., 2016), revealing them as Achilles’s heels that can be exploited in resurgent Trojan horse-style chemical warfare strategies with large membrane-impermeable antibiotics that are desperately needed against multi-drug resistant pathogens with a diderm envelope(Braun et al., 1976; Luna et al., 2020; Negash et al., 2019; Terra et al., 2021). Sideromycins are natural conjugate antibiotics chemically equipped with a siderophore moiety (Lin et al., 2019; Liu et al., 2018) for uptake via TBDRs that recognize iron-bound molecules(Braun, 2009; Luna et al., 2020; Pugsley et al., 1987), yet other metallophore-importing TBDRs may also be suitable targets.

When TBDRs recognize their substrate, import typically occurs by triggering the displacement of the N-terminal plug domain that obstructs the channel (Fujita et al., 2019; Killmann et al., 1993; Ratliff et al., 2022; Shultis et al., 2006). Opening (ungating) of the channel and concomitant substrate import is powered by the PMF-dependent IM complex composed of ExbBD-TonB in which TonB interacts directly with the TBDR and the ExbBD complex is responsible for the mechano-chemical coupling of the proton flux across the IM, leading to structural changes in TonB and indirectly to the rearrangement of the plug within the TBDR (Braun, 2009; Hickman et al., 2017; Ratliff et al., 2022). TBDR-encoding genes are particularly abundant in the genomes of oligotrophic bacteria such as the α-proteobacterium *Caulobacter crescentus* (Nierman et al., 2001), presumably to facilitate the acquisition of a wide range of nutrients from environments where they are scarce. While an abundance of TBDR may seemingly maximize the acquisition of important nutrients, it is paramount for bacteria to regulate the composition of their TBDR-laden OM appropriately such to maximize the efficiency of acquisition while minimizing the risk of mistakenly importing toxic compounds.

Sensory two-component systems (TCSs) are known to control OM remodeling in response to envelope stresses(Konovalova et al., 2017; Laloux and Collet, 2017; MacRitchie et al., 2008; Meng et al., 2021). These TCSs typically include a sensory histidine kinase (HK) in the IM that detects a cue in the extracellular/periplasmic space and then subsequently signals this change by way of a phosphoryl transfer reaction to a cognate cytoplasmic response regulator (RR) which then modulates gene expression via a DNA-binding output domain (Gao and Stock, 2010; MacRitchie et al., 2008; Ruiz and Silhavy, 2005; Stock et al., 1990).

*C. crescentus* could potentially load the OM with 67 TBDR orthologs (Nierman et al., 2001), but most are not expressed under standard growth in rich medium (PYE, Figure S1) (Siwach et al., 2021) and the underlying regulatory networks controlling this TBDR repertoire remain underexplored. The ChvT TBDR is highly expressed in PYE and confers sensitivity to the large hydrophilic glycopeptide antibiotic vancomycin (VNC)(Vallet et al., 2020) (Figure 1A). While VNC typically inhibits cell wall synthesis in Gram-positive (monoderm) bacteria with a minimal inhibitory concentration (MIC) in the microgram/milliliter (μg/mL) range, it is kept at bay by the OM of Gram-negative pathogens such as *Escherichia coli* and *Pseudomonas aeruginosa* (Krishnamoorthy et al., 2016; Ruiz et al., 2006). It is unusual that VNC can also inhibit growth of *C. crescentus* at low MIC (20 μg/mL) and genetic analyses revealed the TBDR ChvT as a determinant promoting VNC susceptibility (Frohlich et al., 2018; Vallet et al., 2020). By contrast, the ChvGI (<u>c</u>hromosomal <u>v</u>irulence factor) TCS promotes VNC in *C. crescentu*s via negative regulation of ChvT (Frohlich et al., 2018; Quintero-Yanes et al., 2022; Vallet et al., 2020) (Figure 1A). ChvGI comprises the ChvG HK and the ChvI DNA-binding RR and while recent studies provided evidence that low pH and sucrose stress trigger ChvGI in *C. crescentus*, other evidence suggests that it can also be induced late in the DNA damage response or after long term growth in the presence of cell wall-targeting antibiotics at sub-MIC levels (Frohlich et al., 2018; Quintero-Yanes et al., 2022; Stein et al., 2021; Vallet et al., 2020). The ChvGI pathway was originally discovered through its role in virulence gene expression and acid stress adaptation in rhizobial lineages (Chen et al., 2008; Li et al., 2002; Lu et al., 2012; Reed et al., 1991), but recent experiments suggested that ChvGI activation in rhizobia after 24 hours of growth in the presence of cell wall-targeting antibiotics (Williams et al., 2022). The *C. crescentus* ChvI targets includes >160 promoters (Stein et al., 2021), many of which drive expression of envelope (stress) proteins (Quintero-Yanes et al., 2022). One of the ChvI targets is the promoter of *chvR* (P*_chvR_*), controlling transcription of a small regulatory RNA acting as post-transcriptional silencer of ChvT translation (Frohlich et al., 2018) (Figure 1A), but a recent study reported that the *chvT* promoter is itself a target of ChvI (Quintero-Yanes et al., 2022).

**Figure 1:**
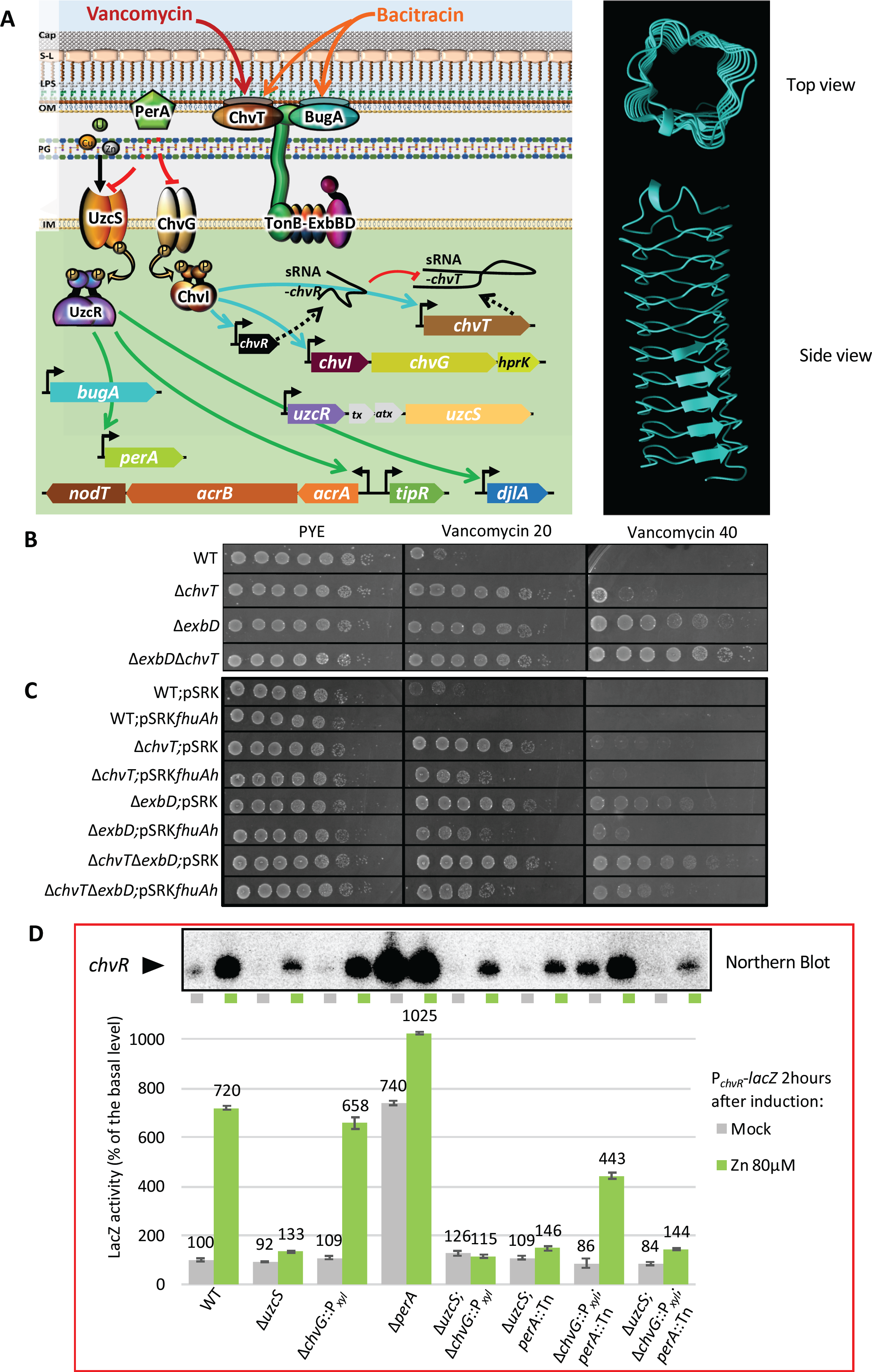
Model of PerA-dependent signalling and identification of *perA*. A) (Left) Proposed model of the TCSs-dependent regulation by PerA across the *C. crescentus* envelope consisting of capsule (cap), S-layer (S-L), an outer membrane (OM) habouring lipopolysaccharide (LPS) in the outer leaflet, cell wall layer (peptidoglycan, PG) and the inner or cytoplasmic membrane (IM). (Right) Alpha-fold predicted three-dimensional structure for the pentapeptide repeat domain (without signal sequence or C terminal domain) in version 1.4 (Jumper et al., 2021; Varadi et al., 2021). B) and C) Efficiency-of-plating (EOP) assays by tenfold serial dilutions of overnight cultures of WT and mutants on PYE plates with or without vancomycin 20 µg/mL or 40 µg/mL grown for 48 hours at 30°C. All plates containing strains harboring the pSRK-Gm or pSRK*fhuAh* plasmid contain IPTG (isopropyl β-D-1-thiogalactopyranoside) at a final concentration of 0.5 mM). C) β-galactosidase activity quantification using the pP*_chvR_-lacZ* promoter probe plasmid. Inductions was performed in exponential phase cultures induced for two hours with ZnSO_4_ (80µM) during growth at 30°C in PYE. All levels are indicated in percentage of expression regarding the basal level of the *WT* (NA1000) in the uninduced condition. The inset shows *chvR* mRNA levels as determined by Northern blotting in the same strains. The blots contain total RNA extracted from cells with (green) and without (grey) zinc induction and the blots were probed for *chvR* mRNA. The full blot is shown in Figure S3.

Interestingly, P*_chvR_* is also a direct target of UzcR, the DNA-binding response regulator component of the UzcSR TCS (Park et al., 2017) (Figure 1A). Binding of UzcR to P*_chvR_* is upregulated in the presence of excess zinc (Zn), copper (Cu) or uranium (U) (Park et al., 2017) and it has therefore been suggested that the HK UzcS, directly or indirectly, senses these metals by an unknown mechanism. Upon Zn induction UzcR also binds promoters of genes encoding TBDRs and other OM components of trans-envelope systems such as the tripartite multidrug efflux pump AcrAB-NodT in which NodT serves as the typical OM portal of RND-type efflux pumps (Blair and Piddock, 2009; Kirkpatrick and Viollier, 2014). Thus, Zn stress may promote remodelling of the OM proteome via UzcSR and the role of *chvR* in silencing ChvT implies a regulatory network promoting OM proteome reprogramming by TBDRs replacing ChvT and perhaps other TBDRs.

Our studies below report our genetic quest to uncover factors controlling OM proteome remodelling using the regulatory network controlling P*_chvR_* as entry point. Our screen unearthed the enigmatic OM protein PerA (Cao et al., 2012; García-Bayona et al., 2019), a pentapeptide repeat protein predicted to fold into a quadrilateral beta-helix (Figure 1A), as major stress regulator controlling OM remodelling and antibiotic uptake. We discovered that induction of TBDRs and other proteins via activation of two intertwined TCSs, UzcSR and ChvGI, coordinated by PerA. Moreover, (Zn) stress induction leads to the proteolytic destabilization of PerA and loss of PerA causes ectopic expression of the hitherto uncharacterized TBDR BugA (CCNA_00224) that confers sensitivity to large antibiotics that impede cell wall synthesis. In summary, our work provides evidence that PerA adopts a central role in orchestrating the OM proteome and the uptake of large in *C. crescentus* through a TCS network regulating TBDRs and other envelope components.

## RESULTS

### Vancomycin sensitivity is conferred by a TBDR system

Loss of the polarity factor TipN renders (Δ*tipN*) cells hypersensitive to VNC for unknown reasons, a sensitivity serving in genetic selections to reveal the TBDR ChvT as a determinant of VNC sensitivity in Δ*tipN* cells (Kirkpatrick and Viollier, 2014). However, it is still unknown which ExbBD and TonB components energize the ChvT trans-envelope system for the uptake of VNC. Predicting that these elusive components would also promote VNC sensitivity, we mapped non-essential VNC-sensitivity determinants in NA1000 (wild-type, *WT)* cells in a single stroke using transposon (Tn) insertion site sequencing (Tn-Seq). To this end, we selected for *himar1* Tn mutants on rich agar medium (PYE) containing VNC (20 μg/mL, VNC^20^) and upon deep-sequence analyses identified three genes as the most frequently disrupted by the Tn: *CCNA_00324* (encoding an ExbD family protein), CCNA_03052 (encoding a putative acetyltransferase) and CCNA_03108 (encoding ChvT) (Figure S2). As expected, the transcripts of all three genes are expressed during growth in PYE, with *chvT* being the most abundant transcript (Figure S1). To confirm the Tn-Seq results, we engineered mutant strains harboring an in-frame deletion in each of two of these genes (Δ*chvT* and Δ*exbD*) and generated the double Δ*chvT*Δ*exbD* mutant to explore a potential cumulative effect. We tested the efficiency of plating (EOP) of the resulting mutant strains on plates containing VNC^20^ or VNC^40^ (Figure 1B). Loss of ExbD or ChvT provided a clear benefit to colony formation on VNC^20^ compared to *WT* cells, with an EOP exceeding that of *WT* cells by four orders of magnitude. Additionally, the EOP of Δ*exbD* Δ*chvT* double mutant cells appeared slightly increased and denser compared to the EOP of Δ*exbD* cells on VNC^40^ plates. In support of this result, when we used Δ*chvT* cells as a starting point for Tn mutagenesis to selecting for *himar1* Tn mutants that grow on VNC^40^ plates, one out of the three clones isolated had a Tn insertion in *exbD*, indicating that loss of ExbD has an additive effect. As loss ChvT or ExbD likely renders the OM of *C. crescentus* less permeable to VNC, we reversed this effect by transforming our strains with a plasmid expressing the ungated TBDR-derived FhuA (also known as TonA) (Braun et al., 1976) channel encoded by the engineered hyperpore *fhuAh* gene variant (Mohammad et al., 2011). As seen in Figure 1C, hyperpore expression partially restored VNC^20^ sensitivity to Δ*chvT* and Δ*exbD* single and double mutant cells. The reduction in EOP is even more pronounced on VNC^40^, revealing a higher protection of Δ*chvT* Δ*exbD* double mutant cells compared to Δ*exbD* single mutant cells by two orders of magnitude. This finding suggests that ExbD controls ChvT function and implicates (an)other TDBR(s) in using this ExbBD-TonB system to import VNC.

### PerA negatively regulates transcription of the sRNA *chvR*

To identify cues and molecular pathways promoting a switch of TBDRs in the OM, we used the transcriptional regulation of *chvR* as a genetic entry point to find regulatory mutants that silence ChvT. The unusual aspect of *chvR* is that it enables a *forward* selection for mutants in which ChvT expression is *down-regulated* (Frohlich et al., 2018) owing to hyperactivation of P*_chvR_*. To this end, we engineered a transcriptional reporter in which P*_chvR_* drives expression of the NptII neomycin phosphotransferase (P*_chvR_-nptII*) that confers resistance to kanamycin. Next, we transformed *WT* cells with the P*_chvR_* -*nptII* reporter plasmid (pP*_chvR_* -*nptII*) and then mutagenized the resulting cells with a *himar1* Tn (conferring gentamycin resistance), selecting for clones with elevated P*_chvR_* activity on (PYE) plates containing kanamycin (15 μg/mL) and gentamycin (1.5 μg/mL). This selection yielded two clones, each harboring a Tn insertion in the enigmatic *perA* gene (*CCNA_01968)*. *perA* is predicted to encode an unusual 183-residue protein with an N-terminal secretion signal sequence followed by a stretch of penta-peptide repeats that are predicted to fold into a quadrilateral β-helix (Figure 1A). While the function of *perA* remains poorly understood, the PerA protein has been localized to the OM (Cao et al., 2012; García-Bayona et al., 2019). Mutations in *perA* and *chvT* surfaced among other mutations in a selection for mutants that survive growth inhibition by the contact-dependent bacteriocin CdzC/D (García-Bayona et al., 2019). Interestingly, *perA* has also been classified as an essential gene in high-density Tn-Seq analysis of the *C. crescentus* genome (Christen et al., 2011). In these experiments viable Tn mutants were selected on plates supplemented with nalidixic acid (NAL), a quinolone antibiotic that targets the DNA gyrase A subunit (GyrA). NAL is often used to counter-select *E. coli* donor cells delivering the Tn to *C. crescentus* recipients by intergeneric conjugation (Christen et al., 2011). While *C. crescentus* cells are resistant to NAL by means of a naturally occurring polymorphism in GyrA, NAL also results in the massive induction of the RND-type multi-drug efflux pump AcrAB-NodT, a tripartite trans-envelope assemblage that can also expel compounds including NAL (Kirkpatrick and Viollier, 2014). Thus, *perA* may be only essential for viability when plates are supplemented with NAL and our work described below supports this view.

Since our pP*_chvR_* -*nptII* selection regime implicated *perA* as a negative regulator of P*_chvR_*, we wanted to quantify P*_chvR_* activity in the absence of PerA. To this end, we first engineered cells with an in-frame deletion in *perA* (Δ*perA*) and transformed them with a low-copy LacZ-based promoter probe plasmid, pP*_chvR_* -*lacZ*, for measurements of P*_chvR_* activity using *lacZ*-encoded β-galactosidase (Figure 1D). We also transformed pP*_chvR_-lacZ* into *WT* cells for comparison and determined that inactivation of *perA* in *WT* results in a 7-fold increase in P*_chvR_* activity (740%). Because the ChvI RR binds P*_chvR_* and has been implicated in P*_chvR_* regulation(Frohlich et al., 2018; Quintero-Yanes et al., 2022), we then engineered a Δ*chvG*::P*_xyl_* strain (henceforth Δ*chvG*) that lacks a functional HK ChvG and simultaneously harbors the xylose-inducible P*_xyl_* promoter downstream of the ChvG coding sequence to maintain expression of the distal genes in the operon. This Δ*chvG* mutation does not bring about a major change in P*_chvR_-lacZ* activity in *WT* cells, but when Δ*chvG* is introduced into the *perA* mutant background, the overactivation of P*_chvR_* -*lacZ* is lost (Figure 1D, lower panel). We also conducted Northern blotting with total RNA preparation from these strains, probing for the chvR mRNA and obtaining matching results (Figure 1D, upper panel, Figure S3). We conclude that loss of the enigmatic OM protein PerA leads to hyperactivation of P*_chvR_* and this hyperactivity requires the ChvGI TCS.

### Loss of PerA renders sensitive to induction of the AcrAB-NodT efflux pump

Since we encountered no difficulties in constructing *perA* mutant strains, either by generating Δ*perA* cells using the standard double recombination (SacB-mediated counterselection strategy) or by backcrossing the *perA*::Tn mutation using ΦCr30-mediated generalized transduction, we speculated that *perA* mutant cells must be sensitive to NAL^20^ (20 μg/mL NAL) and therefore would not surface in the Tn-Seq experiments by Christen *et al*. (Christen et al., 2011). NAL sensitivity of pleotropic envelope mutants is not unprecedented in *C. crescentus*, as evidenced by the finding that cells lacking the polarity factor TipN or the RNA polymerase associated factor TrcR are also NAL-sensitive (Delaby et al., 2021; Kirkpatrick and Viollier, 2014). To probe for a possible NAL-sensitivity of *perA* mutant cells, we conducted EOP assays of *WT* and Δ*perA* cells on PYE with or without NAL^20^ and observed a strong reduction in EOP of five orders of magnitude in cells lacking PerA versus *WT* cells. By contrast, no reduction in EOP was evident for *WT* cells plated on media with or without NAL^20^ (Figure 2A).

**Figure 2:**
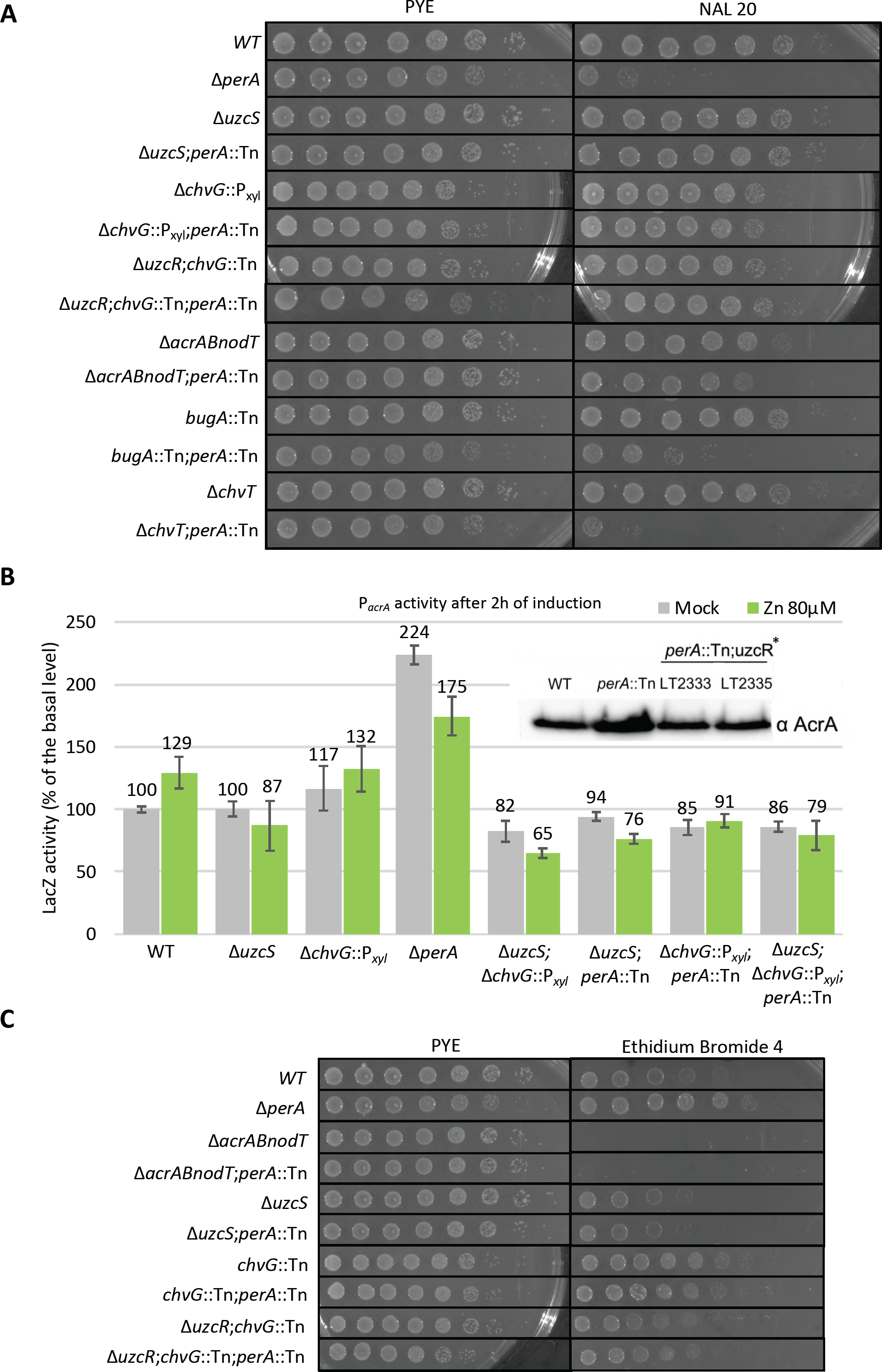
Genetic interplay of *perA, acrAB-nodT* and *uzcR*. A) EOP assays of WT and mutants on PYE plates with or without nalidixic acid 20µg/mL, NAL20) after 48 hours of growth at 30°C. B) β-galactosidase activity measurements using the pP*_acrA_-lacZ* promoter plasmid in various *WT* and mutant cells in exponential growth, before and after ZnSO_4_ (80µM) induction at 30°C in PYE. All levels are indicated in percentage of expression regarding the basal level of the uninduced *WT*. (Inset) Immunoblot analysis using antibodies to AcrA to probe extracts of NA1000 (WT) and *perA* mutant cells that had been separated by SDS-PAGE and blotted. Extracts were made from cells in exponential growing phase in PYE at 30°C. C) EOP assays to assess the efflux capacity of strains grown on PYE plates with or without ethidium bromide (4 µg/mL) for 48 hours at 30°C.

In complementary experiments, we conducted a negative screen for NAL^20^-sensitive Tn mutants in *C. crescentus* by replica-plating an arrayed library of Tn mutants maintained in individual wells of 96-well plates on plates with and without NAL^20^ (Huitema et al., 2006), described in Supplementary Materials and Methods). After mapping the Tn insertions in mutants that grow poorly on NAL^20^ plates, we re-discovered Tn insertions in *perA, tipN* and *trcR* among other genes. This re-isolated *perA*::Tn mutant was designated NAS-4 (<u>NA</u>L-<u>S</u>ensitive, henceforth *perA*::Tn^NAS4^) and harbors a (Hyper Mu) Tn encoding kanamycin resistance. We backcrossed the *perA*::Tn^NAS4^ mutation into *WT* cells and found that the resulting mutant also has a substantially reduced EOP on NAL^20^ plates, akin to Δ*perA* cells. Moreover, we found that the *perA*::Tn^NAS4^ cells also over-activate P*_chvR_* (as determined by LacZ-based quantification with pP*_chvR_* -*lacZ,* Figure S4). Importantly, both these *perA* mutant defects were corrected upon expressing a functional version of PerA *in trans* (Figure S4) from pMT335 (Thanbichler et al., 2007). Thus, PerA silences P*_chvR_* and it confers protection towards NAL^20^.

Since exposure of *C. crescentus* cells to NAL causes a massive transcriptional induction of the RND-type efflux pump AcrAB-NodT (Kirkpatrick and Viollier, 2014), we considered the possibility that *perA* mutant cells fail to grow on NAL^20^ due to a toxic induction of AcrAB-NodT that may destabilize an already fragile or stressed envelope. If this was the case, then deletion the *acrAB-nodT* operon should attenuate the NAL^20^ sensitivity of *perA* mutant cells. Indeed, the EOP on NAL^20^ increased by four orders of magnitude when the *perA*::Tn^NAS4^ was introduced into a mutant background lacking AcrAB-NodT (Δ*acrAB-nodT*, Figure 2A). However, the EOP was not restored to the *WT* level (still exhibiting a two-fold higher EOP), suggesting that the *perA* mutation also acts through other determinants.

To genetically disentangle the pleiotropic phenotypes associated with loss of PerA, we conducted suppressor (pseudo-reversion) analyses, selecting for *perA* mutant that have become NAL^20^-resistant. Such mutants appeared readily upon spreading *perA*::Tn on NAL^20^ plates, suggesting that loss-of-function mutations arise. Indeed, genome sequencing of three such NAL^20^-resistant mutants (designated LT2333, LT2335 and LT2336) revealed different mutations in *uzcR*, encoding the RR UzcR (Figure 1A) (Park et al., 2017). The nonsense mutations in the *uzcR* allele of strain LT2333 is due to a CA dinucleotide insertion at codon 46, inducing a frameshift that terminates 36 codons later, while a C nucleotide insertion at codon 81 in *uzcR* of LT2336 induces termination four codons later. By contrast, mutant LT2335 encodes a missense mutation (CèA) at codon 188 resulting in the UzcR(T188N) variant.

To independently confirm that loss of the UzcRS TCS abrogates the NAL^20^ sensitivity of *perA* mutants, we transduced *perA*::Tn^NAS4^ into Δ*uzcS* cells (lacking the UzcS HK) and found the resulting double mutant cells to have an EOP on NAL^20^ comparable to *WT* cells (Figure 2A). Complementation analysis of Δ*uzcS* strains with a synthetic *uzcR* gene encoding a variant with a double HA-tag at the N-terminus (2xHA-UczR, Figure 2A) restored NAL^20^ sensitivity to the Δ*uzcS*;*perA*::Tn mutant strains when expressed from the *xylX* locus, showing that all three *uzcR* point mutants are recessive (henceforth collectively referred to as *uzcR*\*).

Since activated UzcR binds the *acrAB-nodT* promoter (P*_acrA_*), we conducted promoter probe assays with pP*_acrA_* -*lacZ* in *perA*::Tn single mutant and the three *perA*::Tn *uzcR\** double mutant strains (LT2333, LT23335 and LT2336, Figure S4). These measurements revealed that disruption of *perA* caused a three to four-fold increase in P*_acrA_* -*lacZ* activity, whereas this increase was lost in all *perA*::Tn *uzcR*\* double mutant strains. A similar attenuation of 76% and 91% was observed in *perA*::Tn^NAS4^ Δ*uzcS* and to 91% *perA*::Tn^NAS4^ Δ*chvG* double mutant cells, respectively, compared to the *perA* single mutant (Figure 2B). Moreover, immunoblots using polyclonal antibodies to AcrA, revealed an increase in in the steady-state levels of AcrA in *perA*::Tn cells, whereas *perA*::Tn *uzcR*\* double mutant cells contained *WT* levels of AcrA (Figure 2B, inset). To show that loss of PerA also resulted in an increase in AcrAB-NodT activity, we probed for AcrAB-NodT-mediated efflux of the toxic DNA intercalant ethidium bromide (EtBr) by EOP assays on plates harboring EtBr (4 μg/mL, Figure 2C). Whereas Δ*acrAB-nodT* cells cannot grow on EtBr^4^ plates, *perA*::Tn cells exhibited an *increase* in EOP of approximately three orders of magnitude relative to *WT* cells that was dependent on AcrAB-NodT. By contrast, the EOP of Δ*uzcS* single mutant or *perA*::Tn^NAS4^ Δ*uzcS* double mutant cells is similar or slightly reduced to that of *WT* cells.

We conclude that loss of PerA causes an UzcR-dependent increase of AcrAB-NodT expression that protects cells from EtBr. The elevated basal expression of AcrAB-NodT may in turn also magnify overproduction of AcrAB-NodT in the presence of NAL, compared to the AcrAB-NodT levels in NAL-induced *WT* (see P*_acrA_* -*lacZ* activity assays, Figure S4). However, it is also possible that *perA* mutants are simply more susceptible to insertion of extra AcrAB-NodT in the envelope owing to an elevated OM proteome density or other envelope problems.

### Stress-induced destabilization of PerA activates two intertwined TCSs

Previous chromatin immunoprecipitation-deep sequencing (ChIP-Seq) analyses of *WT* cells revealed that UzcR binds P*_acrA_* and P*_chvR,_* after exposure to Zn stress (Park et al., 2017). In support of this result, our promoter probe assays showed that P*_chvR_-lacZ* and P*_acrA_-lacZ* are rapidly induced by Zn in an UzcS-dependent manner (Figure 1D and 2B) along with the *chvR* transcript ias determined by Northern blotting (Figure 1D). While loss of PerA induces both reporters, again in an UzcS-dependent manner, the addition of Zn did not magnify the response for P*_acrA_-lacZ* in *perA* cells and, conversely, constitutive over-expression of PerA from the vanillate-inducible P*_van_* promoter on pMT335 attenuated the inducibility by Zn (Figure S4). In the case of P*_chvR_-lacZ*, LacZ activity was elevated by the addition of Zn in *perA* mutant cells. However, this response to Zn was attenuated in Δ*chvG perA*::Tn double mutant cells compared to *perA* single mutant cells. Thus, our findings show that modulating PerA levels can affect Zn induction bidirectionally and they suggest that loss of PerA triggers two TCSs: UzcSR and ChvGI.

To demonstrate that the DNA-binding RRs UzcR and ChvI are indeed both more active in *perA* mutant cells versus *WT* cells, we conducted ChIP-Seq experiments with N-terminally HA (hemagglutinin)-tagged versions of UzcR (HA-UzcR) or ChvI (HA-ChvI) expressed from the xylose-inducible promoter at the *xylX* locus in cells with or without PerA. We then determined the PerA-dependent targets based on comparison of genomic binding sites (peaks) detected by the MACS software, using a four-fold enrichment cut-off in the peaks observed in *perA* mutants and versus the isogenic parent (Figure 3A). These ChIP-Seq analyses revealed increased binding of HA-UzcR and HA-ChvI at 134 and 130 target promoters in the absence PerA, respectively (Fig ure3B). From these ChIP-Seq data sets for HA-UzcR and HA-ChvI, we defined an overlapping set of 30 PerA-dependent promoters targeted by both RRs such as the promoters of *chvR*, the TBDR-encoding genes *chvT, CCNA_00974, CCNA_01051* and *CCNA_03096*, the *exbBD-tonB1* operon (*CCNA_02421-CCNA_02419*), and the promoter of *perA*. Thus, UzcR and ChvI target largely separate regulons through distinct target motifs (as determined by MEME-derived sequence logos, Figure 3B) (Park et al., 2017; Quintero-Yanes et al., 2022), yet they are functionally intertwined through negative regulation by PerA.

**Figure 3:**
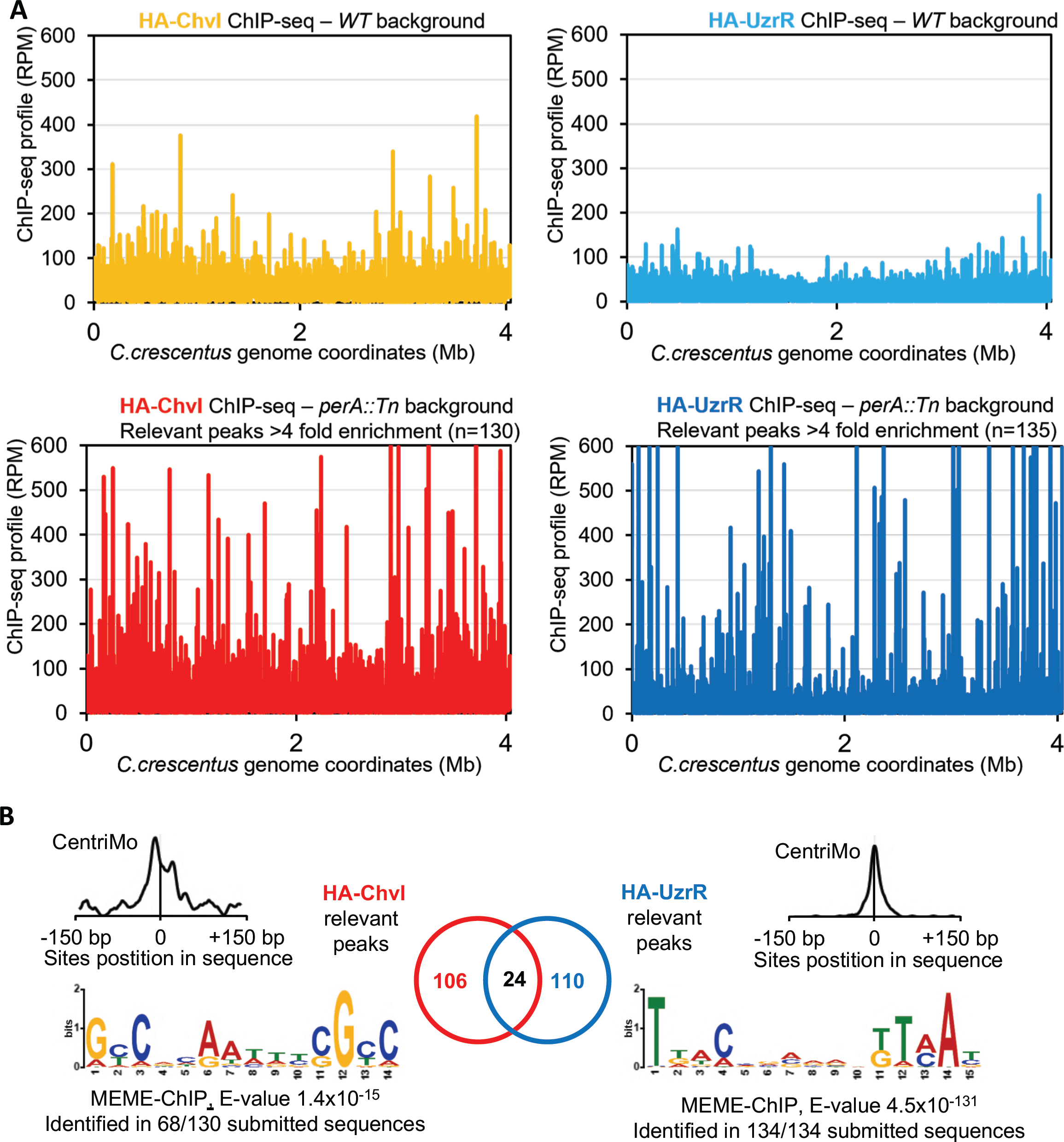
PerA-dependent promoter occupancy of ChvI and UzcR. A) Graphical representation of all HA-UzcR or HA-ChvI binding sites on the chromosome of *C. crescentus* (the X axis is a linear illustration of the genome starting at the *ori*) when 4 times more represented compared to the total input as determined by ChIP-Seq analyses. The Y axis represents the abundance of sequence reads expressed ad Reads Per Million (RPM). B) Venn diagram representing the HA-UzcR and HA-ChvI regulons and the overlapping controlled genes. The MEME-derived sequence logos show the consensus sequence of binding of HA-UzcR and HA-ChvI obtained from respectively 130 and 134 binding sites detected by ChIP-Seq analysis. The CentriMo plots show that the binding sites overlap with the summit of each ChIP-Seq peak, centered between 150 bp upstream or downstream of the peak.

Knowing that UczSR and ChvGI are inducible by Zn and sucrose stress, respectively (Park et al., 2017; Quintero-Yanes et al., 2022), we asked whether these stress conditions can modulate the abundance of PerA. To this end, we engineered *perA*::Tn cells expressing a PerA variant with a C-terminal HA tag from the *xylX* locus (*perA*::Tn *xylX*::*perA-HA*). Expression of PerA-HA in *perA*::Tn cells restores NAL^20^ resistance, indicating that Per-HA is functional (Figure S4). We exposed exponentially growing *perA*::Tn *xylX*::*perA-HA* cells to 80 μM ZnSO_4,_ MgSO_4_ or CuSO_4_ for two hours and monitored PerA-HA abundance by immunoblotting using monoclonal antibodies to HA (Figure 4A). While the abundance of PerA-HA in the presence of MgSO_4_ was similar to that of the untreated control condition, PerA-HA was no longer detectable after ZnSO_4_ exposure and barely detectable in the presence of CuSO_4_. Because acid stress (pH 5.5) has also been reported to induce the *chvR* transcript via ChvGI (Frohlich et al., 2018; Vallet et al., 2020), we tested whether low pH or sucrose stress also destabilizes PerA-HA. Indeed, two hours after applying pH or sucrose stress, cells no longer contained detectable PerA-HA by immunoblotting using anti-HA antibodies (Figure 4A).

**Figure 4:**
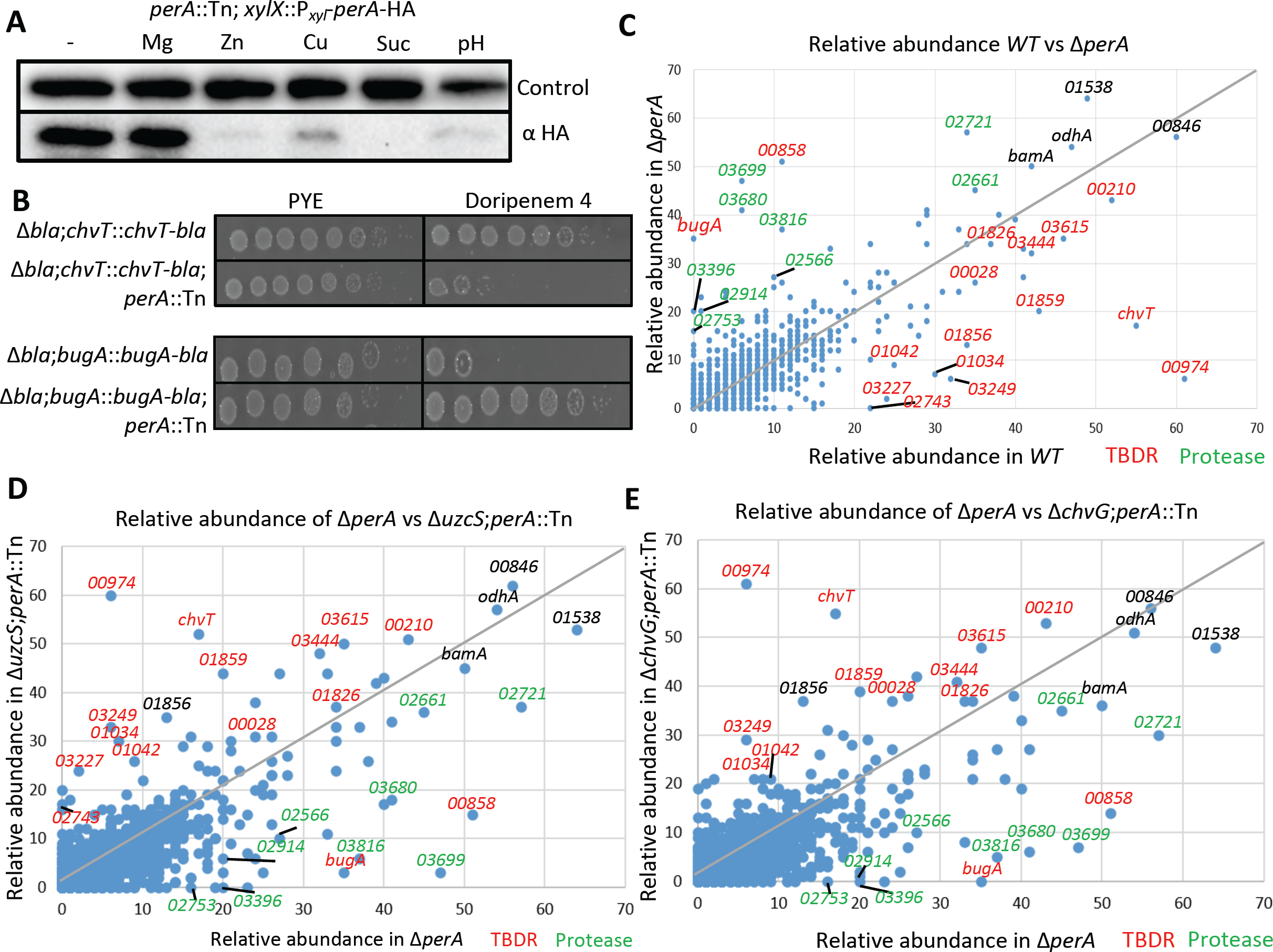
PerA-dependent TBDR remodeling of the OM proteome. A) Immunoblot analysis performed with anti-HA antibodies on the *perA*::Tn; *xylX*::P*_xyl-_perA-*HA strain cells grown with xylose 0,03% for 4h, with and without induction by MgSO_4_, ZnSO_4_, CuSO_4_ (all at 80µM), sucrose 6% or at pH5.5. for two hours. As loading control and immunoblot is shown with antibodies to CCNA_00164. B) EOP assays of strains expressing a ChvT or BugA variant fused to the Bla β-lactamase (ChvT-Bla or BugA-Bla) expressed from the *chvT* or *bugA* promoter at the endogenous locus. EOP was on PYE plates with or without doripenem 4 µg/mL. C), D) and E) Scatter plot representation of the relative abundance of outer membrane enriched protein fractions from WT *versus* the Δ*perA* (C), or Δ*perA versus* Δ*uzcS;*perA::Tn^NAS4^ (D), or Δ*perA versus* Δ*chvG;*perA::Tn^Gm^ cells (E) as determined by LC-MS/MS. The TBDRs are indicated in red while the predicted proteases are colored in green.

These experiments support the conclusion that stress-induced destabilization of PerA triggers two partially converging envelope stress TCSs: ChvGI and UzcSR. Importantly, both of these RRs target a myriad of promoters encoding different TBDRs or other components of trans-envelope assemblies, suggesting that PerA acts as central organizer of the *C. crescentus* OM and/or envelope.

### PerA controls the OM proteome and antibiotic uptake by TBDRs

To determine if *perA* mutants remodel their OM proteome and TBDR repertoire relative to *WT* cells, we conducted OM proteomic analyses by Liquid Chromatography–Mass Spectrometry (LC-MS/MS) of OM protein-enriched samples obtained by differential solubilization of *WT* and Δ*perA* cellular extracts (Figure 4C). The peptides detected by LC-MS/MS revealed an abundance of TBDRs in the OM of *WT* cells, including ChvT and the top 20 highly expressed TBDRs known from RNA-Seq (Figure S1). Comparison of the OM-enriched proteome sample from *WT* cells to that of Δ*perA* cells revealed a massive downregulation of many of these TBDRs in Δ*perA* cells. The downregulation of TBDRs ranged from 50% to more than 98%, in some cases leaving a residual relative abundance of only 2.3% (CCNA_00974) or 1.4% (ChvT/CCNA_03108) in Δ*perA* cells relative to *WT* cells.

By contrast to this down-regulation of some TBDRs, others were massively upregulated in the OM-enriched fraction from *perA* mutant cells versus *WT* cells. For example, we detected a 700-fold induction of the TBDR CCNA_00224 (see below) and a 10-fold induction of the TBDR CCNA_00858 (whose promoters are both targets of UczR (Park et al., 2017)). We also found many predicted extra-cytoplasmic proteases/peptidases to be highly upregulated in the OM-enrichment from Δ*perA* cells compared to *WT*, for example CCNA_03396, CCNA_03699, CCNA_02914 and CCNA_02933. Most of the upregulated proteins are expressed from promoters that are UzcR or ChvI targets. Importantly, the changes in abundance in *perA* mutant cells were reversed as determined from the analysis of OM-enriched samples from the Δ*uzcS perA*::Tn and Δ*chvG perA*::Tn double mutants by LC-MS/MS (Figure 4D and 4E).

To confirm that the massive downregulation of the ChvT protein is not a result of a detection or fractionation problem, we used a genetic approach to show the strong down-regulation of ChvT in *perA*::Tn cells. In these experiments, we used a *C. crescentus* mutant (Δ*bla*) deleted for the gene encoding a metallo-β-lactamase (Docquier et al., 2002; West et al., 2002) to engineer a derivative expressing a ChvT-Bla translational fusion from the *chvT* locus (Δ*bla chvT*::*chvT-bla)*. We then transduced the *perA*::Tn mutation into these cells by transduction and conducted EOP assays on plates with or without the doripenem (4 μg/mL), a β-lactam antibiotic that is enzymatically cleaved by Bla (Docquier et al., 2002). These EOP assays revealed that the *perA*::Tn derivative grows poorly on plates with doripenem (Figure 4B) compared to the isogenic reporter strain. Thus, the ChvT-Bla is no longer expressed in the absence of PerA or it is rendered unstable.

*perA* mutant cells should exhibit an increased EOP on VNC compared to *WT* cells, owing to the apparent absence of ChvT. Surprisingly, we observed the opposite: a drastic decrease in EOP of Δ*perA* cells versus *WT* cells on VAN^7.5^ plates (Figure 5A). We also found a comparable reduction in EOP assays on plates containing bacitracin (3 μg/mL, BAC^3^) for *perA* mutant cells relative to *WT* cells. BAC is a large soluble peptide antibiotic that also targets cell wall synthesis and is excluded by a functional OM, akin to VNC. To illuminate the basis for the BAC sensitivity of Δ*perA* mutant cells, we selected for mutants that grow on BAC^2^ plates following *himar1* Tn mutagenesis. This selection revealed two mutants, each with a Tn insertion in the TBDR-encoding gene *CCNA_00224* (henceforth called *bugA*, for <u>B</u>AC <u>u</u>ptake <u>g</u>ene A). EOP assays revealed that the *bugA*::Tn mutation not only reversed the growth defect of *perA* mutant cells on BAC^3^ plates, but also restored EOP on VAN^7.5^ plates (Figure 5A).

**Figure 5:**
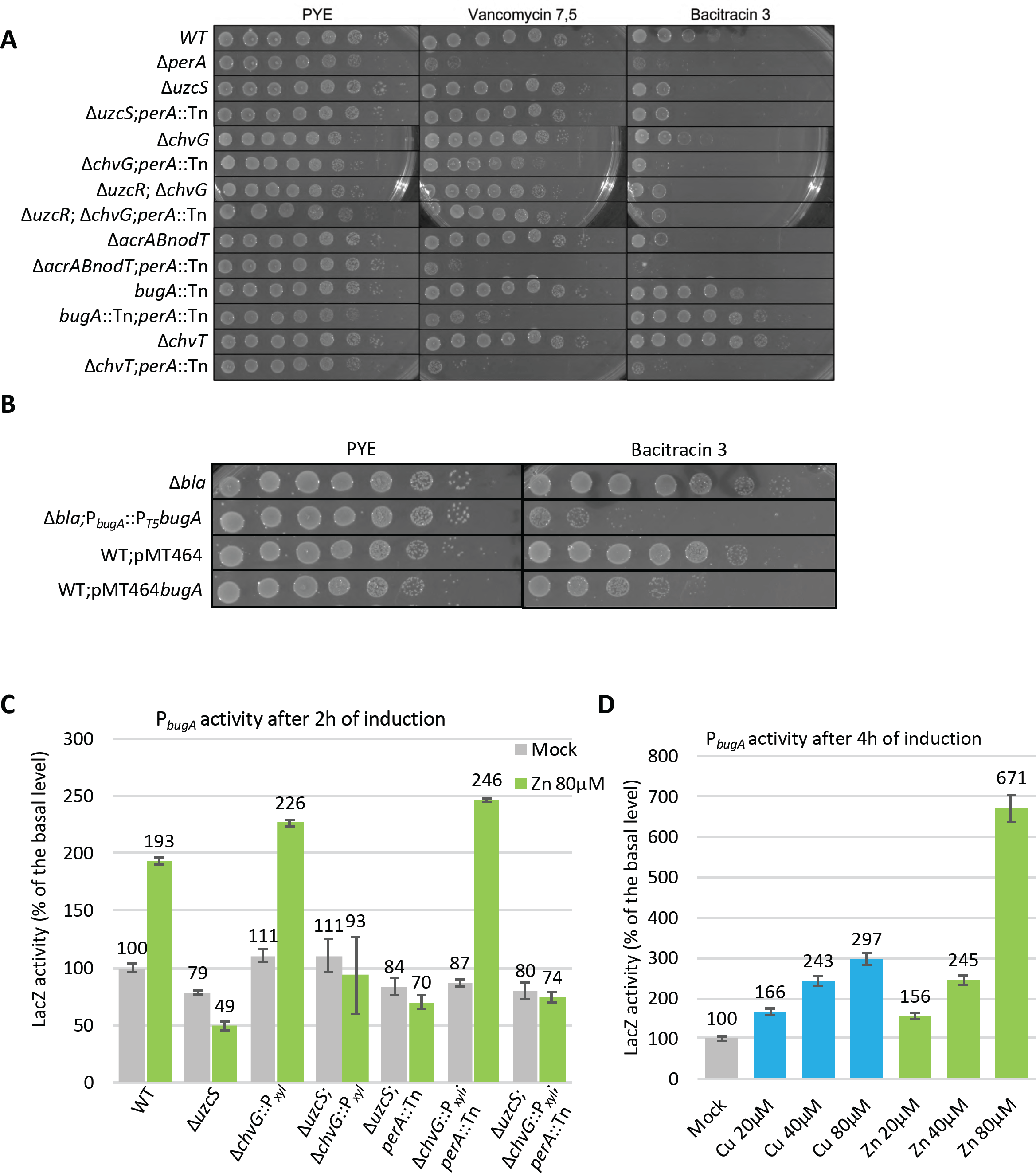
Ectopic expression of BugA confers sensitivity to bacitracin. A) and B) EOP assays of *WT* and *perA* mutant cells on PYE with vancomycin 7,5, 20 or 40 µg/mL or bacitracin 3 µg/mL grown for 48 hours at 30°C. B) and D) β-galactosidase assay using the pP*_bugA_-lacZ* reporter plasmid in WT cells. Inductions was performed two hours (C) or four hours (D) after induction with CuSO4 or ZnSO4 (80µM). All levels are indicated relative to the 100% expression level of the uninduced state.

Remarkably, the BugA TBDR is among the most abundant proteins detected by LC-MS/MS in the OM-enriched fraction prepared from Δ*perA* cells versus *WT* cells, and BugA is no longer detectable in the OM-enriched fractions from Δ*uzcS perA*::Tn and Δ*chvG perA*::Tn cells. Moreover, LC-MS/MS analysis of OM-enriched fractions prepared from WT after two hours of induction with Zn, revealed a strong upregulation of the BugA protein (Figure S5). Since the *bugA* promoter (P*_bugA_*) is among the top targets of UczR (Figure 3A), we confirmed using a pP*_bugA_-lacZ* promoter probe plasmid that P*_bugA_* fires upon Zn or Cu induction (Figure 5D). This activation no longer occurs in cells harboring a Δ*uzcS* mutation or a Δ*uzcS perA*::Tn double mutation (Figure 5C). As Δ*perA* transformants with the pP*_bugA_-lacZ* plasmid were unstable (presumably due to elevated LacZ levels that inhibit *C. crescentus* growth), we were unable to quantify the extent of P_bugA_ hyperactivity in the absence of PerA alone. However, we engineered an Δ*bla bugA*::*bugA-bla* strain that was subsequently transduced with the *perA*::Tn mutation. EOP assays on plates harboring doripenem (4 μg/mL) revealed that the *perA*::Tn mutation rendered these cells much more resistant to doripenem compared to the isogenic parent (Figure 4B). Since loss of PerA results in strong upregulation (or stabilization) of BugA-Bla, we conclude that elevated BugA expression in *perA* mutant cells promotes BAC uptake, whereas BugA is poorly expressed in *WT* cells that are therefore less susceptible to BAC.

To test whether ectopic expression of BugA is sufficient to confer BAC susceptibility to *WT* cells, we engineered a strain in which bugA is expressed from the strong coli phage T5 promoter at the chromosomal *bugA* locus (*bugA*::T5-*bugA*, Figure 5B). This strain exhibited a markedly reduced EOP on BAC^3^ plates compared to the isogeneic parent. Additionally, we engineered a replicative plasmid (pMT464-BugA) expressing BugA from the *xylX* promoter and also observed that pMT464-BugA imparts a substantial reduction in the EOP of *WT* cells on plates BAC^3^, compared to the empty vector (pMT464, Figure 5B). Knowing that BugA also reduces the EOP of *perA* mutant cells on VNC^7.5^, we next determined whether pMT464-BugA also reduces the EOP of Δ*chvT* cells on plates containing VNC^20^ or VNC^40^ compared to empty the empty vector (Figure S6). Indeed, we noted a reduction in EOP in the presence of pMT464-BugA, a reduction that also occurs in Δ*exbD* single mutant and in Δ*chvT* Δ*exbD* double mutant cells (Figure S6), indicating that BugA can function independently of ExbD (CCNA_00324), likely using more than one ExbBD-TonB systems indistinctly.

In summary, our result show that *bugA* is necessary and sufficient to confer sensitivity to VNC and BAC in *C. crescentus*, explaining how loss of PerA results in sensitivity to large cell wall targeting antibiotics, despite the absence of ChvT. In a physiological context our work supports the model that stress-induced destabilization of PerA changes the OM proteome and TBDR-mediated antibiotic uptake through induction of two conserved TCSs.

## DISCUSSION

The decision when to dynamically remodel the OM proteome with TBDRs is a critical regulatory juncture for bacterial survival. In the light of potentially fatal trade-off between food acquisition versus (suicidal) antibiotic import, it is paramount to illuminate the underlying mechanisms of induction as these could potentially lead to new (combined) options to treat infections by Gram-negative bacteria (Luna et al., 2020). The BugA TBDR confers sensitivity to VNC and BAC that bind cell wall precursors, indicating that these antibiotics possess related structural determinants to engage this TBDR. However, BAC and VNC likely also engage the TBDR ChvT, perhaps less efficiently since expression of BugA substantially increases the sensitivity to these antibiotics, regardless of whether cells express ChvT or not (Figure 5A and 5B). Since zinc-stressed cells execute a TBDR switch from ChvT to BugA, it may endow the OM with a TBDR having improved uptake kinetics of cell wall (related) products. In light of the fact that VNC and BAC are Zn-binding antibiotics (Economou et al., 2013; Zarkan et al., 2017; Zarkan et al., 2016), our findings raise the intriguing possibility that VNC and BAC can enter and kill cells akin to metallophore-induced uptake as a “zincophore” Trojan-Horse antibiotic that may prove useful to diderm bacteria under stress conditions. In fact, BAC can also enter *Mycobacterium tuberculosis* (*Mtb*) cells through a multi-solute transporter that also allows the recently discovered gyrase-targeting antibiotic evybactin to traverse the OM (mycomembrane) of *Mtb* (Imai et al., 2022).

OM re-modelling is often coordinated by TCSs in response to stress (Raivio and Silhavy, 1999; Saha et al., 2021). To impede overcrowding and destabilization of the OM, cells may need to reduce the number of pre-existing TBDRs by stopping their synthesis followed by dilution or proteolytic decay as new TBDRs are being inserted. Indeed, the stress inducible regulon characterized here for *C. crescentus* includes newly expressed TBDRs, proteases/peptidases, as well the BamA and BamB components of the β-barrel insertion machine (BAM) for most OM proteins (Hagan et al., 2011). Unexpectedly, our work unveiled a β-helical OM protein serving as stress-induced orchestrator of multiple TCSs. In *Escherichia coli,* the RcsF OM lipoprotein is thought to act as an OM stress-sensor activating a TCS that mitigates imbalances or defects in envelope integrity. RcsF is a BAM substrate that can reach through the BamA lumen to the OM surface (Cho et al., 2014; Konovalova et al., 2014; Laloux and Collet, 2017; Meng et al., 2021; Saha et al., 2021). When BAM function is perturbed for example by OM/envelope deformations and/or the increased abundance of other BAM substrates, less BamA may be available for translocation of RcsF to the cell surface and for insertion of other OM proteins. Since RcsF triggers the Rcs TCS in the periplasm through relief of negative regulation, an imbalance in BAM function and/or alterations of the OM surface may affect RcsF localization or folding, ultimately triggering Rcs (Cho et al., 2014; Tata et al., 2021). However, is also possible that RcsF controls an accessory BAM activity. While OM localization and negative TCS regulation are also hallmarks of the PerA stress responsive system, it remains to be determined whether PerA (that lacks a prototypical lipidation motif such as the one found in RcsF) is a substrate of the *C. crescentus* BAM.

While we found Zn stress as a trigger of the PerA-dependent transcriptional response, yet it is evident that other stresses also destabilize PerA. How such diverse stress responses can affect PerA stability is not clear. In the case of Zn stress, PerA might bind Zn ions directly, resulting in structural deformations that render it susceptible to proteolysis by specific extra-cytoplasmic proteases, for example CtpA-type carboxyl-terminal processing proteases (Sommerfield and Darwin, 2022). Interestingly, there is also precedence for another β-helix protein, GlmU from *Streptococcus pneumoniae*, to bind Zn (Brazel et al., 2022). GlmU is an essential and bifunctional cell wall biosynthetic metalloenzyme in the cytoplasm that synthesizes UDP-linked N-acetyl glucosamine (UDP-GlcNAc) through a globular uridyl-transferase domain in the N-terminal region, but is also possesses an β-helical acetyltransferase domain in the C-terminal region whose activity is impaired when it binds Zn (Brazel et al., 2022).

It also possible that Zn stress affects PerA indirectly through other targets. For example, the addition of Zn ions could displace the calcium and magnesium ions in the LPS or impair cell wall biosynthesis (perhaps via GlmU) to liberate or activate periplasmic proteases or to enable their access to PerA. Mis-metallization by Zn could alter the substrate spectrum of a metal-dependent periplasmic protease that act on PerA. Interestingly, *C. crescentus* Bla is also a zinc-dependent metalloenzyme whose activity could potentially be affected by Zn levels (Docquier et al., 2002) (West et al., 2002).

By analogy to the rhizobial ChvGI system in which a periplasmic regulator, ExoR (Reed et al., 1991), downregulates the orthologous TCS (in an unknown way), a different negative regulator ChvGI TCS has long been predicted for the *C. crescentus* which does not encode a detectable ExoR ortholog (Nierman et al., 2001). ExoR is also degraded in response to acidification (Lu et al., 2012), but it is a soluble α-helical protein located in the periplasm. Surprisingly, PerA not only regulates ChvGI, but also the UzcSR TCS which is also encoded in rhizobial genomes, begging the question whether this TCS is also engaged by acid stress in rhizobia. Acid stress can also affect the envelope, specifically cell wall biosynthetic activities in different ways. PBP1a is inhibited by low pH (Mueller et al., 2019) and a number of other cell wall factors with roles in low pH have been reviewed (Mueller and Levin, 2020), indicating that common perturbances between pH and Zn stress are conceivable.

*perA* and *chvT* both surfaced (among others) in a previous screen for resistance to the bacteriocin CdzC/D (García-Bayona et al., 2019). Since it is known that TBDRs can also serve as entry and uptake portals for bacteriocins and we have shown that *perA* mutants down-regulate ChvT, this TBDR may serves as an entry point for CdzC/D. If true, then *perA* mutants may simply be resistant to this bacteriocin simply because they downregulate ChvT and possibly additional TBDRs that bind and import the bacteriocin.

Owing to their potential utility as narrow range antimicrobials that avoid perturbance of the native microbiome, TBDRs and similar uptake systems hold great potential in highly targeted Trojan horse style chemical warfare against Gram-negative and other diderm pathogens using natural or semisynthetic metal-bound antibiotics (Wencewicz and Miller, 2018). Finally, we highlight a situation where induction of the genetically-integrated RND-efflux pump AcrAB-NodT is lethal to bacterial cells with an altered OM proteome, raising the possibility that antibiotics targeting the BAM complex(Kaur et al., 2021; Miller et al., 2022) might exhibit strong lethal synergies with those that induce these efflux pumps. Loss of PerA elevates AcrAB-NodT-mediated efflux of toxic molecules, for example towards ethidium bromide, it also renders cells sensitive to massive overexpression of this efflux pump for reasons that remain to be determined.

## EXPERIMENTAL PROCEDURES

### Strains and growth conditions

*Caulobacter crescentus* NA1000 and derivatives were cultivated in peptone-yeast extract (PYE) at 30 °C incubator. Liquid cultures were grown on rotary wheel and ФCr30-mediated generalized transductions were done as described (Ely, 1991). *E. coli* was grown in LB (Luria Bertani) with antibiotics at 30 °C under agitation in media 0.3 mM meso-diaminopimelic acid (mDAP) was supplemented to enable growth for WM3064 derivatives (Saltikov and Newman, 2003). Plates contained 1.5% agar medium assay, and soft agar overlays were done by mixing ¼ solid medium with ¾ of liquid PYE. Liquid and solid media used for *C. crescentus* were supplemented with antibiotics at the indicated concentrations (in µg/mL). When mentioned, xylose is used at concentrations of 0.03%, IPTG at 0.5μM and vanillate at 100μM for inducing the expression of corresponding promoters. Transposon delivery plasmids were pHVPV414 (Viollier et al., 2004) or pMAr2xT7(Liberati et al., 2006). Strains and plasmids are summarized in Supplementary Table S1.

### Efficiency of plating (EOP) assays

Petri dishes were prepared with the mentioned antibiotic(s). ON cultures were normalized at OD_600_ = 0.5. Tenfold serial dilutions were performed by transferring 20µl of the culture in 180µl of fresh media in 96 wells plates. This step was repeated until the 8 dilutions were done. Five μL of each dilution was dropped on the selected Petri dishes and incubated for 48 hours at 30°C.

### Genome-wide transposon mutagenesis coupled to deep-sequencing (Tn-Seq)

Overnight cultures of the *E. coli* donor cells S17-1 λ*pir* strain containing the *himar1*-delivery plasmid pHPV414 with a kanamycin resistance cassette (Viollier et al., 2004), and overnight cultures of the *C. crescentus* recipient cells were mixed at a ratio of 250 µL of the *E. coli* donor cells and 1 mL of *C. crescentus* recipient cells. After centrifugation, cells were resuspended in 40 µL PYE and incubated for 5h at 30°C on PYE agar for mating. Cells were plated either on PYE containing kanamycin 20 µg/mL and NAL^20^ (control) or PYE with kanamycin 20 µg/mL, NAL^20^ and VNC^20^ incubated at 30°C for 5 days. An average of 100’000 Tn mutants on the former plate and polled, while a few thousand were isolated from the latter plate and pooled. Chromosomal DNA was extracted from the pools using Ready-Lyse Lysozyme (Epicentre Lucigen) and DNAzol (ThermoFischer) according to the manufacturers’ recommendations. This harvested DNA was used to generate a Tn-Seq library and submitted to Illumina HiSeq 2500 sequencer (Fasteris, Geneva, Switzerland). The Tn insertion specific reads were sequenced using the *himar1*-based Tn-seq primer (5’-AGACCGGGGACTTATCAGCCAACCTGTTA-3’) to create the single-end sequence reads (50 bp) which were mapped against the genome of the reference genome of *Caulobacter* crescentus NA1000 (NC_011916) (Nierman et al., 2001). Data was processed using the Galaxy software (https://usegalaxy.org/) and SeqMonk software (http://www.bioinformatics.babraham.ac.uk/projects/seqmonk/) to build sequence read profiles. Using SeqMonk, the genome of *Caulobacter* was divided into 50 bp probes, and a calculated value that represents a normalized read number per million has been determine for every probe. A ratio of the reads obtained in the control condition versus the vancomycin treatment was calculated for each 50 bp position. Tn-Seq reads sequencing and alignment statistics are summarized in Supplementary Table S2.

### Chromatin Immuno Precipitation coupled to deep Sequencing (ChIP-Seq) analyses

Cultures of exponentially growing (OD660nm of 0.5, 80 ml per sample of culture in PYE supplemented with xylose 0.3%) *C. crescentus* cultures Δ*uzcR*; *xylX*::P*_xyl_-H*A-*uzcR, the perA mutant derivative* Δ*uzcR*; *xylX*::P*_xyl_*-HA-*uzcR; perA::Tn* (Gm^R^), as well as Δ*bla*; *xylX*::P*_xyl_-HA-chvI* and the *perA* mutant derivative Δ*bla*; *xylX*::P*_xyl_*_-_*HA-chvI; perA*::*Tn* (Gm^R^) were respectively supplemented with 10 μM sodium phosphate buffer (pH 7.6) and then treated with formaldehyde (1% final concentration) at room temperature for 10 minutes to achieve crosslinking. Subsequently, the cultures were incubated for an additional 30 minutes on ice and washed three times in phosphate buffered saline (PBS, pH 7.4). The resulting cell pellets were stored at −80°C. After resuspension of the cells in TES buffer (10 mM Tris-HCl pH 7.5, 1 mM EDTA, 100 mM NaCl) containing 10 mM of DTT, the cell resuspensions were incubated in the presence of Ready-Lyse lysozyme solution (Epicentre, Madison, WI) for 10 minutes at 37°C, according to the manufacturer’s instructions. Lysates were sonicated (Bioruptor Pico) at 4°C using 15 bursts of 30 seconds to shear DNA fragments to an average length of 0.3–0.5 kbp and cleared by centrifugation at 14,000 rpm for 2 minutes at 4°C. The volume of the lysates was then adjusted (relative to the protein concentration) to 1 mL using ChIP buffer (0.01% SDS, 1.1% Triton X-84 100, 1.2 mM EDTA, 16.7 mM Tris-HCl [pH 8.1], 167 mM NaCl) containing protease inhibitors (Roche) and pre-cleared with 80 μl of Protein-A agarose (Roche, www.roche.com) and 100 μg BSA. Five percent of the pre-cleared lysates were kept as total input samples (negative control samples). The rest of the pre-cleared lysates was then incubated overnight at 4°C with monoclonal rabbit Anti-HA Tag antibodies (Millipore, clone 114-2C-7) at a 1:400 dilution. Immuno-complexes were captured by incubation with Protein-A agarose beads (pre-saturated with BSA) during a 2 hour period at 4°C and then, washed with low salt washing buffer (0.1% SDS, 1% Triton X-100, 2 mM EDTA, 20 mM Tris-HCl pH 8.1, 150 mM NaCl), with high salt washing buffer (0.1% SDS, 1% Triton X-100, 2 mM EDTA, 20 mM Tris-HCl pH 8.1, 500 mM NaCl), with LiCl washing buffer (0.25 M LiCl, 1% NP-40, 1% deoxycholate, 1 mM EDTA, 10 mM Tris-HCl pH 8.1) and finally twice with TE buffer (10 mM Tris-HCl pH 8.1, 1 mM EDTA). Immuno-complexes were then eluted from the Protein-A beads with two times 250 μL elution buffer (SDS 1%, 0.1 M NaHCO_3_, freshly prepared) and then, just like the total input samples, incubated overnight with 300 mM NaCl at 65°C to reverse the crosslinks. The samples were then treated with 2 μg of Proteinase K for 2 h at 45°C in 40 mM EDTA and 40 mM Tris-HCl (pH 6.5). DNA was extracted using phenol:chloroform:isoamyl alcohol (25:24:1), ethanol-precipitated using 20 μg of glycogen as a carrier and resuspended in 30 μl of DNAse/RNAse free water.

The immunoprecipitated chromatin was used to prepare sample libraries used for deep-sequencing at Fasteris SA (Geneva, Switzerland). ChIP-Seq libraries were prepared using the DNA Sample Prep Kit (Illumina) following manufacturer instructions. Single-end run was performed on an Illumina Next-Generation DNA sequencing instruments (NextSeq High), 50 cycles (for *HA-uzcR*) and 75 cycles (*HA-chvI*), respectively, were performed and yielded several million reads per sequenced sample. The single-end sequence reads stored in FastQ files were mapped against the genome of *C. crescentus* NA1000 (NC_011916.1) using Bowtie2 Version 2.4.5+galaxy1 available on the web-based analysis platform Galaxy (https://usegalaxy.org) to generate the standard genomic position format files (BAM). ChIP-Seq reads sequencing and alignment statistics are summarized in Supplementary Table S3. Then, BAM files were imported into SeqMonk version 1.47.2 (http://www.bioinformatics.babraham.ac.uk/projects/seqmonk/) to build ChIP-Seq normalized sequence read profiles. Briefly, the genome was subdivided into 50 bp, and for every probe, we calculated the number of reads per probe as a function of the total number of reads (per million, using the Read Count Quantitation option). Analysed data illustrated in Figure 3A and B are provided in Supplementary Table S3. Using the web-based analysis platform Galaxy (https://usegalaxy.org), HA-ChvI and 2xHA-UzcR ChIP-Seq peaks were called using MACS2 Version 2.2.7.1+galaxy0 (No broad regions option) relative to the total input DNA samples. The q-value (false discovery rate, FDR) cut-off for called peaks was 0.05. Peaks were rank-ordered according to their fold-enrichment values (Table S2, Peaks with a fold-enrichment values >4 were retained for further analysis). Consensus sequences common to the top 130 HA-ChvI peaks and to the top 134 HA-UzcR peaks (identified in *perA*::*Tn* backgrounds) were respectively identified by scanning peak sequences (+ or - 150 bp relative to the coordinates of the peak summit) for conserved motifs using MEME-ChIP (Version 5.5.0; http://meme-suite.org/)(Bailey et al., 2006).

### β-galactosidase assays

β -galactosidase assays were done using freshly electroporated strains carrying the pLac290 plasmid with transcriptional fusions between indicated promoters and *lacZ*. Briefly, 100µL of cells of (OD_600_nm= 0.3-0.7) were lysed in 30 µL of chloroform and vigorously mixed with Z buffer (60 mM Na2HPO4; 40 mM NaH2PO4; 10 mM KCl and 1 mM MgSO4; pH 7) to obtain a final volume of 800 µL. To begin the reaction 200 µL of ONPG were added (o-nitrophenyl-b-D-galactopyranoside, at 4 mg/mL in 0.1 M potassium phosphate, pH 7). Assays were stopped with 500 µL of 1 M Na_2_CO_3_ when the solution turned light yellow. All assays were done at room temperature. The OD_420_nm of the supernatant and the OD_420_nm of the culture was collected and use to calculate the Miller units as follows: U=(OD_420_*1000)/(OD_600_*t(min)*v(ml)). The standard deviation is represented as error bars. All data are from a minimum of three biological replicates.

### Immunoblots

Strains were grown for 2h at 30°C under constant agitation up to OD_600nm_ between 0.2 to 0.4, prior the addition of the inducer, and the cells were grown for two additional hours. Samples were harvested by centrifugation and resuspended in SDS sample buffer (50 mM Tris-HCl pH 6.8, 2,5 % SDS, 10% glycerol, 1% β-mercaptoethanol, 12.5 mM EDTA, 0.05 % bromophenol blue). Protein samples were analyzed on an SDS–polyacrylamide (37.5:1) gel electrophoresis and blotted on 0.45 µm pore PolyVinyliDenFluoride (PVDF) membranes (Immobilon-P from Sigma Aldrich). Membranes were blocked 2h with 1X Tris-buffered saline (TBS) (50 mM Tris-HCl, 150 mM NaCl [pH 8]) that contain 0.1% Tween-20 and 8% powdered milk. After 1h of incubation, the primary antibodies were used ON at 4°C. The polyclonal antisera to AcrA (1:15000), HA (1:20000) and CCNA_00164 (1:20000) were used(Ardissone et al., 2014; Vallet et al., 2020). The detection of primary antibodies was done using HRP-conjugated donkey anti-rabbit antibody (Jackson ImmunoResearch) with Western Blotting Detection System (Immobilon from Milipore). Images was done on Bio-Rad illuminator (Chemidoc MP, Biorad).

### Cell fractionation and LC-MS/MS analysis

Fifty mL of an exponential culture of *Caulobacter* at OD_600_nm = 0.4 to 0.6 was harvested by centrifugation for 15 min at 8000xg at 4°C. The cells were washed twice with 20 mL cold PBS, then resuspend in 1mL resuspension buffer (20 mM Tris-HCl pH 7.5, 300 mM NaCl, 12500 U Ready-lyse (Epicentre technologies), and one tablet of EDTA-free protease inhibitor cocktail). After 5 minutes of incubation, 10 μL of DNase 1 μg/mL, 5 μL of RNaseA 20 μg/mL were added prior sonication in an ice-water bath (15 cycles 30 seconds ON; 30 seconds OFF). Sonicated samples are centrifuged for 40 minutes at 20’000xg at 4°C, the supernatant was removed. The pellet was resuspended in 300 μL of 20 mM Tris-HCl pH 7.5, 300 mM NaCl, 2% Triton X-100 and incubated at RT for 20 min. Centrifugation was performed for 30 minutes at 20’000xg at 4°C. The final pellet was solubilized in 2X SDS sample buffer and boiled for 10 min, then centrifuge for 2 minutes at 13’000xg. The supernatant was harvested and sent for MS identification at the Taplin Mass Spectrometry Facility at Harvard Medical School (Boston, USA).

### Northern analysis

Northern blots were done using total RNA extracted from *C. crescentus* and probed for *chvR* as previously described(Frohlich et al., 2018).

## ACKNOWLEDEMENTS

We thank Yves Mattenberger for valuable intellectual input and for constructing pSRK-*fhuAh*. We also thank Sean Crosson, Clare Kirkpatrick, Dan Park and Yongqin Jiao for strains, materials and/or discussions. This work was supported by a Swiss National Science Foundation (Grant CRSII5_198737 to P.H.V), the Canton de Genève and the Deutsche Forschungsgemeinschaft (FR 3502/2-1 to K. S. F.). J.C. owes special thanks to Sabine Quindou, Jamy Gourmaud and Frédéric Courant for nurturing an entire generation’s curiosity.

## Legends to Figures

**Figure S1: Expression of TBDRs as determined by RNA-Seq.**

A) Graphical representation of the htseq-count of all the annotated TBDR in *Caulobcater crescentus* NA1000 from (Siwach et al., 2021).

**Figure S2: Vancomycin resistance determinants identified by Tn-Seq.**

A) Plot representation of the Tn-Seq on vancomycin 20 µg/mL with the top nine genes with the higher ratio of Tn insertion and the six lower.

**Figure S3: chvR mRNA levels in various strains before and after zinc induction.**

Northern blots are shown as described in the legend for Figure 1D. This figure shows the full blot and markers.

**Figure S4: Phenotype of cells lacking PerA.**

A) EOP of *perA* mutant complemented with PerAHA or *uzcR* mutant complemented with HA-UzcR on PYE containing or not nalidixic acid 20µg/mL grown for 48 hours at 30°C. Both plates contain 0.03% of xylose.

B) and C) β-galactosidase assay using the pP*_acrA_-lacZ* promoter probe plasmic various mutants performed for two (B) or four hours (C) with MgSO_4_, ZnSO_4_ (80µM) or nalidixic acid 10µg/mL (+) at 30°C in PYE. All levels are indicated in percentage of expression relative to the basal level of the *WT* in the uninduced condition. All experiments done with strains carrying the pMT335 (and derivatives) were done in the presence of vanillate (100µM).

C) EOP of strains carrying the pLac290 empty or expressing the kanamycin resistance (*nptII*) under the control of P*_chvR_* on PYE containing or not kanamycin 30µg/ml grown for 48 hours at 30°C.

**Figure S5: OM proteome analysis by LC-MS/MS in response to zinc induction.**

Scatter plot graphic representing the relative abundancy of proteins from outer membrane enriched samples from the WT strain after 2 hours of growth with or without ZnSO_4_ (80µM). The TBDR are indicated in red while the predicted proteases are in green.

**Figure S6: BugA can act independently of ExbD.**

Kirby-Bauer type antibiograms of *C. crescentus* strains embedded in PYE soft agar. Antibiotic discs were placed on soft agar from top left to bottom right: colistin (50 µg), fosfomycin (50 µg), piperacillin (100 µg), rifampicin (30 µg), moenomycin (5 µg), bacitracin (130 µg), teicoplanin (30 µg), vancomycin (30 µg) and ramoplanin (40 µg). Plates were grown for 24 hours.

